# A representative reference for MRI-based human axon radius assessment using light microscopy

**DOI:** 10.1101/2021.06.03.446739

**Authors:** Laurin Mordhorst, Maria Morozova, Sebastian Papazoglou, Björn Fricke, Jan Malte Oeschger, Thibault Tabarin, Henriette Rusch, Carsten Jäger, Stefan Geyer, Nikolaus Weiskopf, Markus Morawski, Siawoosh Mohammadi

## Abstract

Non-invasive assessment of axon radii via MRI bears great potential for clinical and neuroscience research as it is a main determinant of the neuronal conduction velocity. However, there is a lack of representative histological reference data on the scale of the cross-section of MRI voxels for validating the MRI-visible, effective radius (*r*_eff_). Because the current gold standard stems from neuroanatomical studies designed to estimate the frequency-weighted arithmetic mean radius (*r*_arith_) on small ensembles of axons, it is unsuited to estimate the tail-weighted *r*_eff_. We propose CNN-based segmentation on high-resolution, large-scale light microscopy (lsLM) data to generate a representative reference for *r*_eff_. In a human corpus callosum, we assessed estimation accuracy and bias of *r*_arith_ and *r*_eff_. Furthermore, we investigated whether mapping anatomy-related variation of *r*_arith_ and *r*_eff_ is confounded by low-frequency variation of the image intensity, e.g., due to staining heterogeneity. Finally, we analyzed the potential error due to outstandingly large axons in *r*_eff_. Compared to *r*_arith_, *r*_eff_ was estimated with higher accuracy (normalized-root-mean-square-error of *r*_eff_: 7.2 %; *r*_arith_: 21.5 %) and lower bias (normalized-mean-bias-error of *r*_eff_: *−*1.7 %; *r*_arith_: 16 %). While *r*_arith_ was confounded by variation of the image intensity, variation of *r*_eff_ seemed anatomy-related. The largest axons contributed between 0.9 % and 3 % to *r*_eff_. In conclusion, the proposed method accurately estimates *r*_eff_ at MRI voxel resolution across a human corpus callosum sample. Further investigations are required to assess generalization to brain areas with different axon radii ensembles.

## 1. Introduction

The MRI signal generated by an ensemble of protons probing the local, microscopic environment in human brain tissue contains information about microstructural tissue features such as the axonal radius [1, 2, 3, 4]. The axonal radius is a key to determine neuronal communication in the human brain because it is related to, e.g., the neuronal conduction velocity [5, 6, 7]. The estimation of the axonal radius and other microstructural features via biophysical modeling of the MRI signal [8] is an active area of research because of its potential to partially replace or complement invasive ex-vivo histology with non-invasive, in-vivo histological MRI (hMRI) [9, 10] approaches. However, before these models can be used, they need to be validated against a robust histological reference. [11].

The validation for the MRI-visible, effective radius (*r*_eff_) is currently lacking a robust, histological reference for human brain tissue. Since *r*_eff_ is indicative of large, sparsely occurring axons, i.e., the tail of the axon radii distribution [1, 12, 13], it is essential to accurately capture the tail of the axon radii distribution in large samples to representatively estimate *r*_eff_ for MRI voxels of a human MRI system (1 mm^3^ or larger). For perfusion-fixed rats, validation of *r*_eff_ has been attempted on histological images of the cross-sectional size of ultra-high-resolution MRI voxels (*∼*100 µm^3^) using a preclinical MRI system [1]. However, validation on human brain tissue remains of interest, because it is unclear whether the validation of MRI-based models on rats can be translated to the human brain. As the tail of the axon radii distribution may vary between humans and other mammals [14, 15], *r*_eff_ for humans may be shifted with respect to other species. This shift may be further reinforced by the reduced capability to resolve small axons in human MRI systems when compared to preclinical MRI systems [1, 16]. For human brain, the current gold standard for the validation of *r*_eff_ [1, 2, 21, 22] stems from neuroanatomical studies [17, 18, 19, 20] of small ensembles of axons (100-1000 axons), aiming to evaluate the arithmetic mean radius (*r*_arith_) on manually annotated electron microscopy images (EM). As *r*_arith_ is weighted towards the bulk of axon radii, it can be expected that estimates of *r*_arith_ are less sensitive to the ensemble size as compared to *r*_eff_. For *r*_eff_, however, small-ensemble estimates can strongly under- or overestimate *r*_eff_ [23] of typical MRI voxels, because the tail of the axon radii distribution is insufficiently sampled.

Albeit high-resolution, large-scale light microscopy (lsLM) cannot resolve small axons as accurately as EM, an lsLM-based approach might be appropriate to generate a histological gold standard for the validation of MRI-based radius estimation in human brain tissue. Because of its large field-of-view, covering cross-sections of 1 mm^2^ or larger, lsLM enables the generation of large ensembles of axons including 10^5^ to 10^6^ axons per section and thus allows for more accurate sampling of the tail of the axon radii distribution. Moreover, lsLM has the advantage of being fast, cheap and simple to perform. As the assessment of axon radii on large field-of-view microscopy data renders manual annotation infeasible, automated approaches, e.g., methods based on convolutional neural networks (CNN), are required. So far, CNN-based methods based on large two- or three dimensional scanning or transmission electron microscopy (SEM/TEM) sections have been trained on images of perfusion-fixed mice or rats [24, 25]. However, it is unlikely that the models generated in these studies translate well to immersion-fixed human brain tissue with higher tissue degradation.

In this study, we investigate the potential of lsLM and CNN-based segmentation to map the distribution of axon radii in a human corpus callosum specimen. We quantify the capability of the proposed method to estimate the MRI-visible *r*_eff_ and the *r*_arith_, which is commonly reported in neuroanatomical studies, by evaluating the estimation errors on six lsLM sections. While reference data for the frequency-weighted *r*_arith_ can be generated through manual annotation with reasonable effort, the tail-weighting of *r*_eff_ introduces the necessity to accurately capture the tail of the axon radii distribution and thus investigate larger ensembles of axons than can be realistically annotated. To address this challenge, we follow a three-fold approach to assess the contribution of different axon radii ranges towards *r*_eff_. First, the error due to badly-resolved, small axons is assessed in a cross-microscopy comparison using annotated EM sections from close spatial proximity as a reference. Second, the error due to medium-sized axons, that represent the bulk of the axon ensemble, is assessed using Monte Carlo methods. Third, the error due to large axons, is assessed based on exhaustive annotation of large axons in large field-of-view sections. Additionally, we investigate whether our method is capable of capturing anatomy-related, spatial variation of *r*_arith_ and *r*_eff_ in the presence of low-frequency image intensity variation, e.g., due to staining heterogeneity. Finally, we analyze the potential error due to individual, outstandingly large axons in *r*_eff_.

## 2. Materials and methods

### 2.1. Ensemble mean axon radii

In the following, we define an axon radii ensemble of *B* individual radii *r*_*b*_ with *b ∈* [1, *B*] as

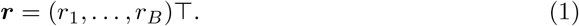

After binning axon radii of this ensemble into *K* bins, the arithmetic mean radius can be defined as

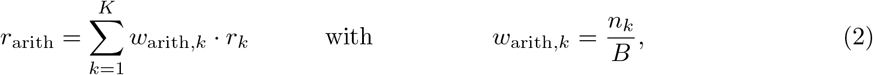

where *n*_*k*_ is the number of axons with radius *r*_*k*_. Analogously, the MRI-visible, effective mean radius [1, 12, 13] can be defined as

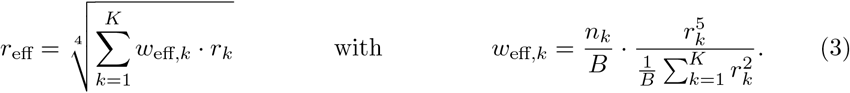

While *r*_arith_ is frequency-weighted (*w*_arith,*k*_), *r*_eff_ is weighted (*w*_eff,*k*_) towards the tail of the axon radii distribution because *w*_eff,*k*_ scales with the fifth power of *r*_*k*_.

In the following, we denote vectors as *bold-faced*, estimations with a *hat* and reference values with a *tilde*. Thus, we denote estimations of individual radii, axon radii ensembles (see Eq. (1)), the arithmetic mean radius (see Eq. (2)) and the effective radius (see Eq. (3)) as 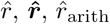, and 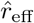; corresponding reference values are denoted as 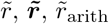 and 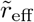.

### 2.2. Data acquisition

Microscopy images from four human white matter samples of four different subjects were acquired: a corpus callosum (CC, male, 74 years, 24 hours postmortem delay, multiple tumors and multi organ failure), a corticospinal tract (CST, female, 89 years, 24 hours postmortem delay, heart failure), an optic chiasm (OC, male, 59 years, 48 hours postmortem delay, multi organ failure) and a sample obtained from the area dorsolateral of the olivary nucleus including the anterolateral system (AS, male, 81 years, 24 hours postmortem delay, multi organ failure). Following standard procedures, blocks were immersion-fixed in 3 % paraformaldehyde and 1 % glutaraldehyde in phosphate-buffered saline at pH 7.4. Then, smaller blocks of 1 to 4 mm edge length were cut, contrasted with osmium tetroxide and uranyl acetate, dehydrated in graded acetones, embedded in Durcupan resin and cut into semi-(*∼*500 nm) and ultra-thin (*∼*50 nm) sections. Semi-thin sections were stained with 1 % toluidine blue.

In total, 13 lsLM images were acquired of semi-thin sections using a Zeiss AxioScan Z1 (objective: 40 *×*, numerical aperture: 0.95, resolution: 0.1112 µm/pixel; resolution limit: 292 nm) (see Table 1). For six regions of the CC sample, lsLM sections and *matching* EM sections were acquired, i.e., sections were cut within 100 µm proximity (see Fig. 1). For the latter EM sections, images were acquired using a Zeiss LEO EM 912 Omega TEM at 80 kV and digital micrographs were obtained with a dual-speed 2K-on-axis CCD camera-based YAG scintillator (TRS-Tröndle, resolution: 0.0043 µm/pixel, resolution limit: 4 nm).

**Table 1:** The dataset of human tissue samples. The following tissue *samples* were investigated: a corpus callosum (CC), a corticospinal tract (CST) an optic chiasm (OC) and an anterolateral system (AS). *Sections* were assigned exclusively to the *training* or *test* dataset.

| Sample | Type | Size [mm<sup>2</sup>] | Sections |  |  |
| --- | --- | --- | --- | --- | --- |
|  |  |  | Total | Training | Test |
| OC | lsLM | 2.49 | 1 | 1 |  |
| CST | lsLM | 12.71 | 1 | 1 |  |
| AS | lsLM | 4.86 | 1 | 1 |  |
| CC | lsLM | 0.34 to 9.16 | 10 | 4 | 6 |
| CC | EM | 0.01 | 6 |  | 6 |

**Figure 1:**
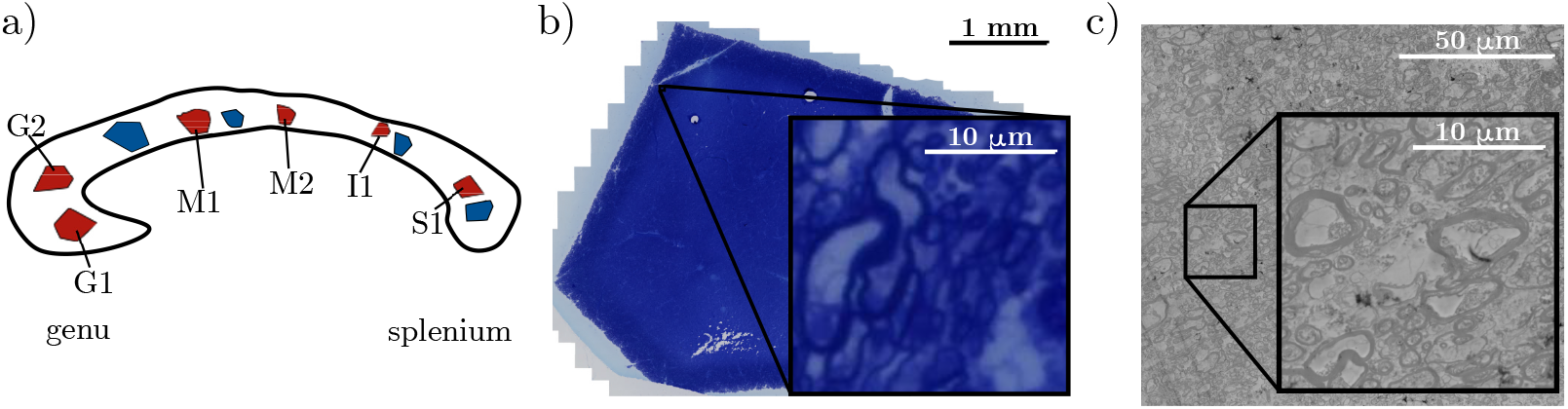
The human corpus callosum sample. The schematic of the sample (a) highlights the regions used for training (blue) and testing (red). For each region, one lsLM section was acquired. For the six test regions (red), *matching* lsLM and EM subsections were acquired: two sections from genu (G1, G2), two sections from midbody (M1, M2) and one section each from isthmus (I1) and splenium (S1). For section G1, the lsLM (b) and its *matching* EM (c) section are depicted as well as examples of subsections that were magnified to cover the same spatial extent (20 *×* 20 µm^2^) at common resolution.

### 2.3. Data annotation

#### 2.3.1. Training dataset

For training of the CNN, we annotated 64 lsLM subsections of similar size (70 *×* 70 µm^2^ to 120 *×* 120 µm^2^) originating from the training sections of the four tissue samples (see Table 1): 46 CC subsections, 4 OC subsections, 4 CST subsections and 10 AS subsections. To cover a wide range of appearance in axon shape and image contrast, some subsections were only partially annotated, i.e., pixels were assigned an ignore label and were not considered during training. As large axons were expected to have particular relevance for *r*_eff_, but occur with low frequency, we assigned higher priority to the annotation of these axons.

#### 2.3.2. Test dataset

We annotated subsections of the *N* = 6 test regions (Fig. 1a) as reference data to assess the error of 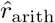 (see Eq. (2)), the error of 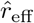 (see Eq. (3)) and the segmentation performance of the CNN in evaluation experiments (see Section 2.6). To account for the different characteristics of these measures (see Section 2.1), we generated separate reference subsections, which differ in the type of microscopy data (EM and lsLM), the subsection size and the annotation scope, i.e., selective and entire annotation of axons of particular size (see Table 2).

**Table 2:**
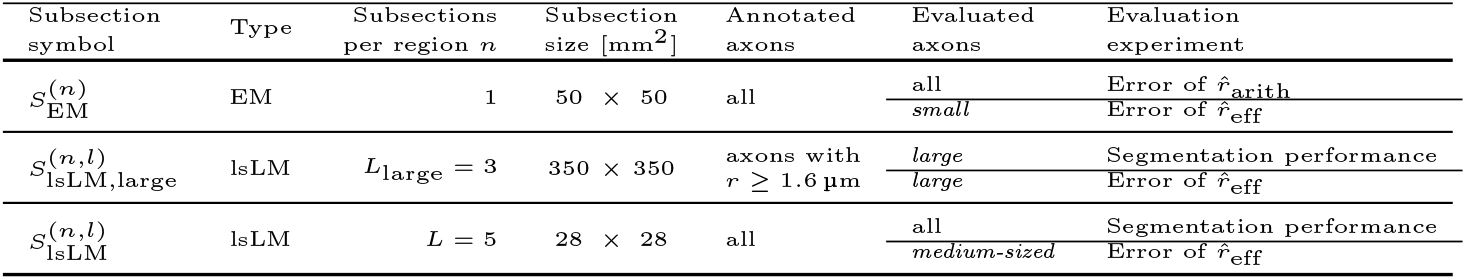
The test dataset. For each of the *N* = 6 test regions, high-resolution, large-scale light microscopy (lsLM) and electron microscopy (EM) subsections were annotated entirely or selectively (column: *annotated axons*). During *evaluation experiments* (see Section 2.6), these annotations were used partially or entirely (column: *evaluated axons*). Axon radii ranges used during *evaluation experiments* are as follows: *small* (*r <* 0.3 µm), *medium-sized* (0.3 *≤*µm *r <* 1.8 µm) and *large* (*r ≥*1.8 µm). *n ∈* [1, *N*] denotes the test region index. *l* denotes the subsection index, which can be between 1 and the number *subsections per region*.

| Subsection symbol | Type | Subsections per region $n$ | Subsection size [$\text{mm}^2$] | Annotated axons | Evaluated axons | Evaluation experiment |
| --- | --- | --- | --- | --- | --- | --- |
| $S_{\text{EM}}^{(n)}$ | EM | 1 | $50 \times 50$ | all | all<br>*small* | Error of $\hat{r}_{\text{arith}}$<br>Error of $\hat{r}_{\text{eff}}$ |
| $S_{\text{lsLM,large}}^{(n,l)}$ | lsLM | $L_{\text{large}} = 3$ | $350 \times 350$ | axons with<br>$r \geq 1.6 \mu\text{m}$ | *large*<br>*large* | Segmentation performance<br>Error of $\hat{r}_{\text{eff}}$ |
| $S_{\text{lsLM}}^{(n,l)}$ | lsLM | $L = 5$ | $28 \times 28$ | all | all<br>*medium-sized* | Segmentation performance<br>Error of $\hat{r}_{\text{eff}}$ |

To assess *r*_arith_, we annotated *matching* EM subsections (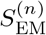 with *n ∈* [1, *N*]; see Table 2). Thereby, we generated references that accurately resolve even *small* (*r <* 0.3 µm) axons below the resolution limit of lsLM, which we expected to have a relevant contribution towards the frequency-weighted *r*_arith_.

As *r*_eff_ is weighted towards the tail of the axon radii distribution (see Eq. (3)), accurate assessment requires sufficient sampling of the tail and thus subsections that cover large ensembles of axons (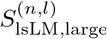 with *n ∈* [1, *N*] and *l ∈* [1, *L*_large_]; see Table 2). Because exhaustive annotation of these large ensembles was infeasible, we annotated only axons with a radius above a certain threshold (*r ≥* 1.6 µm). This threshold was chosen so that 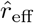 was decreased by 50 % when axons above this threshold were removed from the pooled axon radii ensemble of the corpus callosum lsLM sections evaluated with a prototype of the proposed method. However, for evaluation the threshold had to be adjusted to 1.8 µm (for further explanation see Section 2.6.1). To complement the assessment of 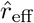 for the error introduced by smaller axons, we used the annotations of *small* axons in *matching* EM subsections *S*_EM_ and the annotations of remaining, *medium-sized* (0.3 µm *≤ r <* 1.8 µm) axons in small lsLM subsections (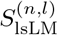 with *n ∈* [1, *N*] and *l ∈* [1, *L*]; see Table 2). The so-obtained split of the axon radii distribution for the assessment of 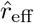 into

1. *small* axons (*r <* 0.3 µm),
2. *medium-sized* axons (0.3 µm *≤ r <* 1.8 µm) and
3. *large* axons (*r ≥* 1.8 µm)

was used throughout different evaluation experiments to assess the contribution of individual axon radii ranges towards 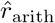 or 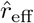.

The segmentation performance of the CNN across the entire range of axon radii was assessed on small subsections 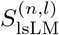, whereas the segmentation performance for *large* axons on 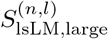.

#### 2.3.3. Raters

Annotation of lsLM subsections, i.e. both training and test subsections, was performed by a total of six raters (M. Morozova, B. Fricke, J.M. Oeschger, S. Papazoglou, T. Tabarin and L. Mordhorst). Each annotated subsection was crosschecked by a second rater. Initially, annotations were carried out in collaboration with two neuroanatomy experts (i.e., M. Morawski and M. Morozova) who were furthermore consulted in case of doubt. EM subsections were annotated by M. Morozova.

### 2.4. Axon radius estimation pipeline

Axon radius estimation was divided into three steps: semantic segmentation, instance segmentation and radius approximation (see Fig. 2). To perform semantic segmentation, i.e., to classify each pixel as either axon, myelin or background, we applied a CNN (see Section 2.5) in a sliding window manner (see Fig. 2a). To identify axon instances from individual pixels, we applied connected-component labeling (see Fig. 2b). Axon radii were approximated as radii of circles with equivalent areas to those of the axon instances (short: circular equivalent) (see Fig. 2c). Axon radii ensembles (see Eq. (1)) were composed from approximated radii. For annotated images, the semantic segmentation step was skipped.

**Figure 2:**
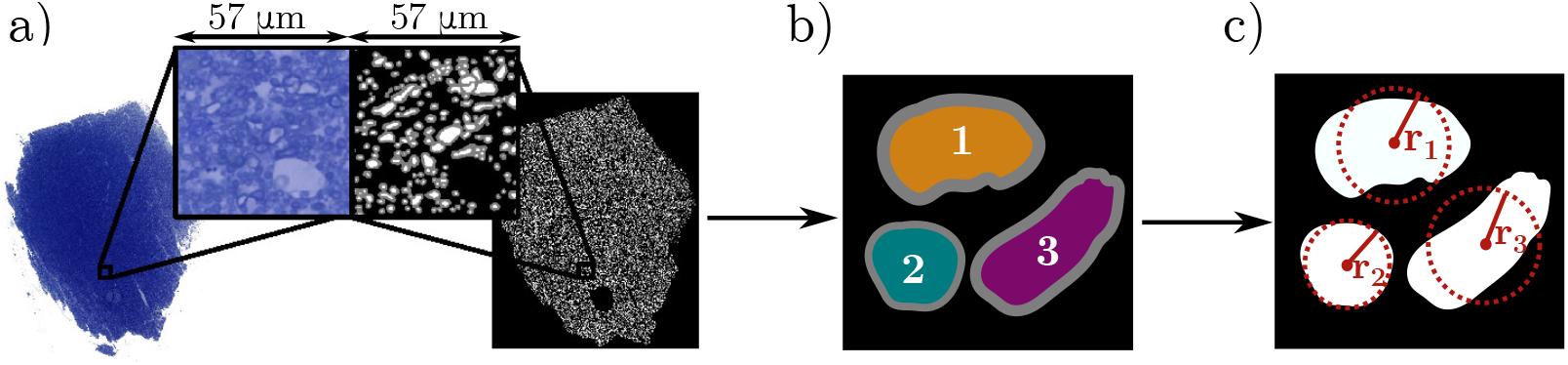
The axon radius estimation pipeline. (a) Pixel-wise classifications as axon, myelin or background were obtained through application of a semantic segmentation network, i.e. a U-Net [26] variant, in a sliding window manner. (b) Subsequently, axon instances were identified through connected component labeling of pixels that were classified as axon. (c) For each axon instance, the radius was estimated as the radius of a circle with equivalent area.

### 2.5. Semantic segmentation network

#### Network Architecture

We used a CNN of the U-Net [26, 27] family with an input size of 512 *×* 512 pixels and four encoder stages. We employed transfer learning: the original encoders were replaced by EfficientNet-B3 [28] encoders that were pretrained on the ImageNet dataset [29].

#### Training and Validation

The training process was split into pseudo-epochs of 150 random patches. Augmentations were employed on-the-fly during training: geometric, brightness and saturation transformations, Gaussian blurring and stain augmentation [30]. Inputs were standardized with respect to the training dataset. We used stochastic gradient descent with Nesterov momentum (0.95), an initial learning rate of 10^*−*2^ and a learning rate decay of *γ* = 0.2 every 50 epochs after initial 100 epochs to minimize a Lov’asz-softmax loss [31]. All weights of the CNN were modified during training. We determined the optimal learning rate, *γ* and the number of epochs in a 4-fold cross-validation with splits conducted at the level of entire subsections. The best model was determined in terms of the averaged dice score for axon and myelin and trained for 200 epochs. We used a framework [32] based on PyTorch [33] to carry out the training procedure.

### 2.6. Evaluation of the performance of the radius estimation pipeline

#### 2.6.1. Performance of the semantic segmentation network

We considered the binary, pixel-wise classification task of discriminating between axon and background, i.e., all non-axon pixels were considered background, to evaluate balanced accuracy, dice, precision and recall. For all *N* test regions, we evaluated all *L* subsections with entirely annotated axons (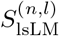 with *n ∈* [1, *N*] and *l ∈* [1, *L*]; see Table 2) and all *L*_large_ subsections selectively annotated for the assessment of *large* axons (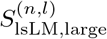 with *n ∈* [1, *N*] and *l ∈* [1, *L*_large_]; see Table 2). To compute the above metrics for 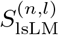, annotations served as an immediate reference for comparison with the prediction. To determine above metrics for 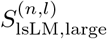, we filtered both annotation and prediction for *large* axons to generate reference and *large*-axon prediction (see Fig. 3). The corresponding aforementioned metrics were then computed using reference and *large*-axon prediction. To summarize each metric over all subsections of all regions for 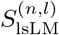 and 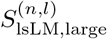, we computed the mean of each metric, respectively.

**Figure 3:**
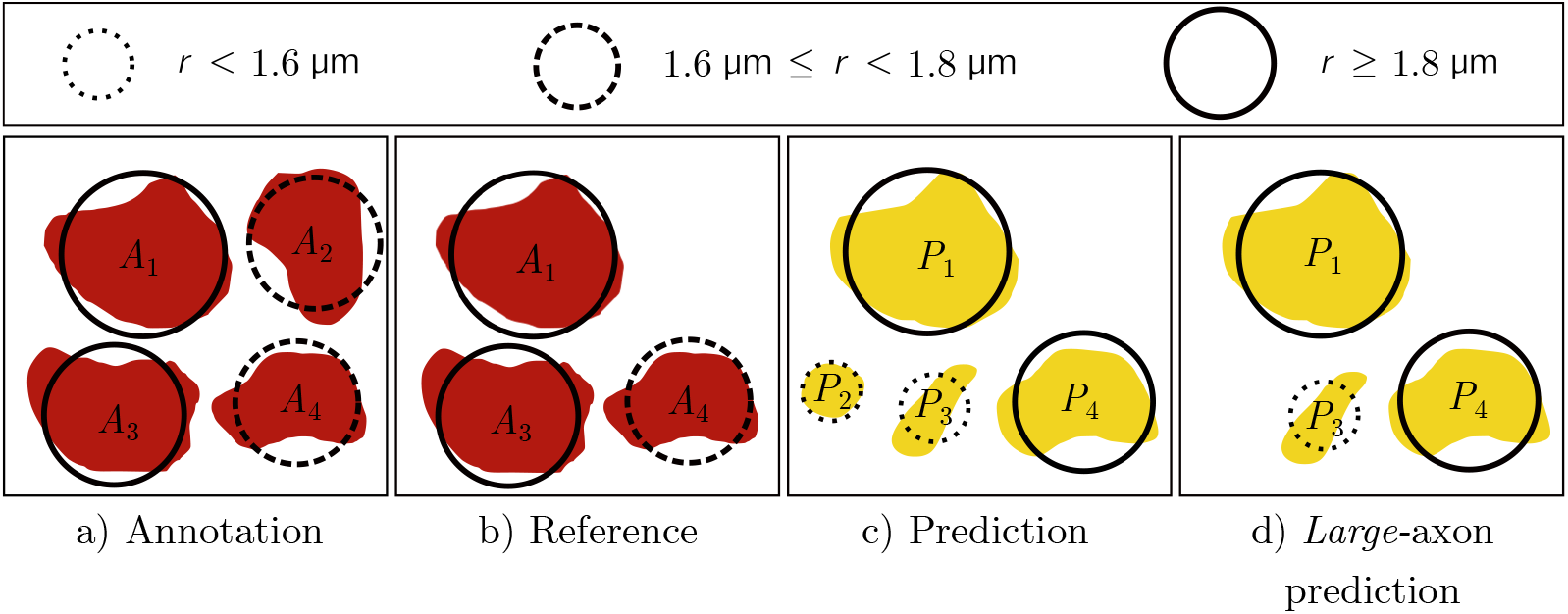
Generation of reference and *large*-axon prediction for one subsection. The annotation (a) was filtered to generate the reference (b). Similarly, the prediction (c) was filtered to generate the *large*-axon prediction (d). For the reference (b), *large* (*r ≥* 1.8 *µ*m) axons were always considered (a-b, *A*_1_ and *A*_3_). Annotated axons with 1.6 µm *≤ r <* 1.8 µm (a, *A*_2_ and *A*_4_) were used as follows: we assumed that predicted, *large* axons (e.g., in c-d, *P*_4_) with no *large*, annotated counterpart but an annotated counterpart with 1.6 µm *≤ r <* 1.8 µm were correctly detected, but marginally oversegmented. Therefore, we considered annotated axons with 1.6 µm *≤ r <* 1.8 µm (a-b, *A*_2_ and *A*_4_) for the reference if a *large*, overlapping axon existed (a-b, *A*_4_) but discarded them otherwise (a, *A*_2_). For the *large*-axon prediction (d), axons were considered if they had *r >*= 1.8 µm (c-d, *P*_1_ and *P*_4_) or if they had the highest pairwise dice similarity (c-d, *P*_1_, *P*_3_ and *P*_4_) to a reference (b) axon. To compute the pairwise dice similarity, only the respective two axons were considered, whereas remaining pixels were considered background. Circles denote circular equivalent axon areas.

#### 2.6.2. Error of 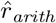

We assessed the error of 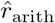 by comparing estimates obtained from lsLM subsections (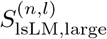 with *n ∈* [1, *N*] and *l ∈* [1, *L*_large_]; see Table 2) against references obtained from annotated EM subsections (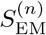; see Table 2). For *l*-th subsection of the *n*-th region, we computed an estimate 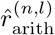 from the predicted axon radii ensemble of 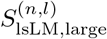 and a reference 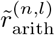 from the EM-based axon radii ensemble 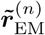 of the *matching*, annotated subsection 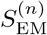. Thereby, references were shared across all *L*_large_ subsections per region, i.e., 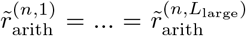. To assess accuracy and bias of 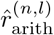 with respect to 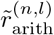 across all *N · L*_large_ subsections of all regions, we computed the normalized-root-mean-square error

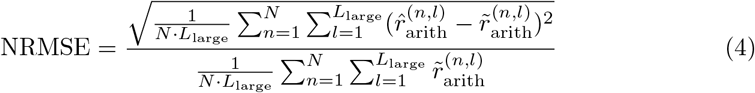

and the normalized-mean-bias-error

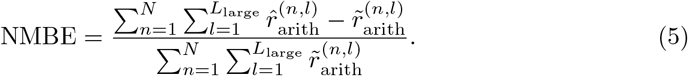

#### 2.6.3. Error of 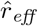

The error of 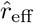 (see Eq. (3)) was assessed on large lsLM subsections (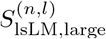 with *n ∈* [1, *N*] and *l ∈* [1, *L*_large_]; see Table 2). To distinctively assess the error due to axons in particular axon radii ranges (see Section 2.3.2), we compared axon radii ensembles with erroneous radii in the particular range against reference values.

For the *l*-th subsection of the *n*-th region, we generated a reference axon radii ensemble 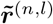 and erroneous axon radii ensembles: one each for the *small* and *large* axon radii range (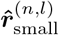 and 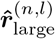); and multiple (*M* = 500) each for the *medium-sized* and entire axon radii range, of which 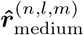 and 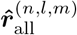 denote the *m*-th (*m ∈* [1, *M*]) erroneous axon radii ensemble (see Fig. 4). The generation of the aforementioned ensembles is detailed in the following paragraphs.

**Figure 4:**
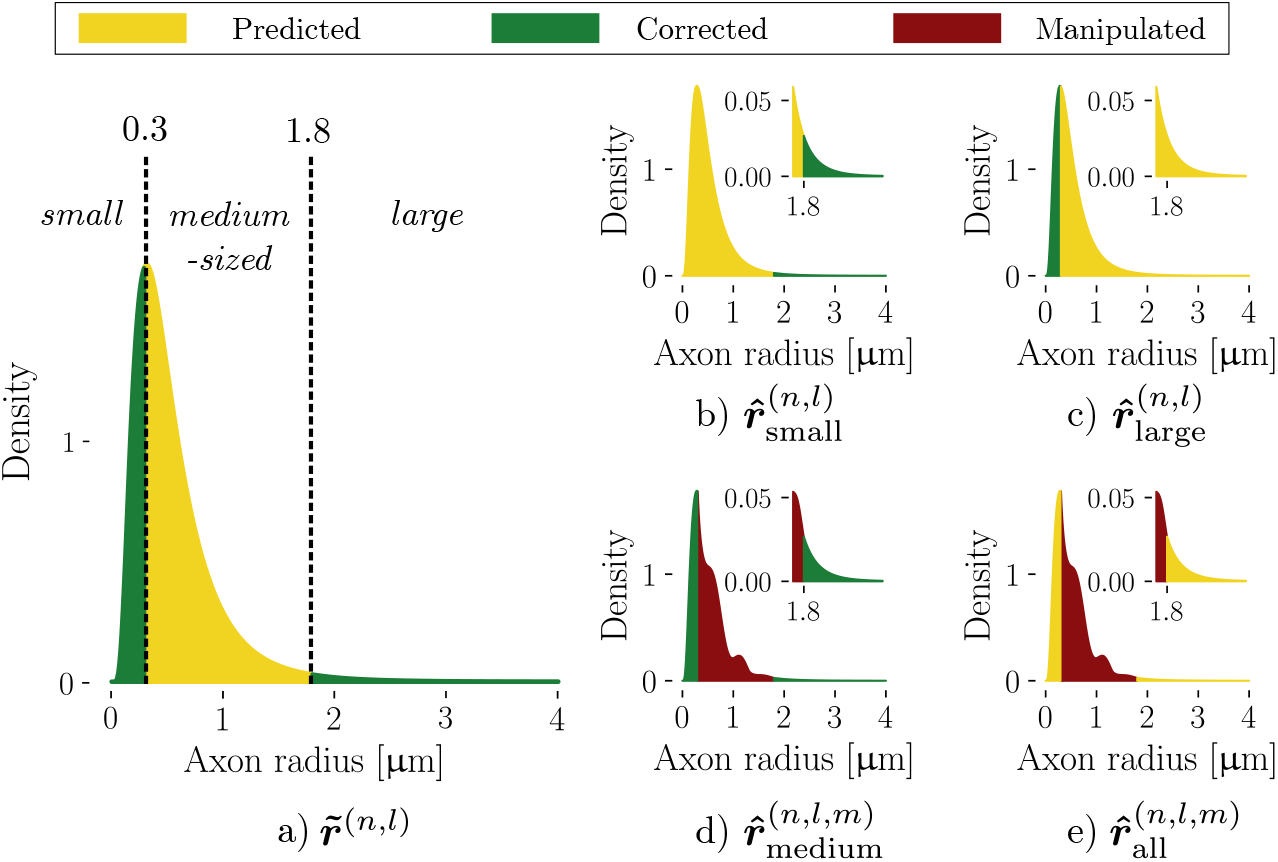
Generation of axon radii ensembles for the error assessment of 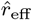. For one subsection (*l* of region *n*), the reference axon radii ensemble (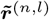; a) was based on the predicted (yellow) axon radii ensemble 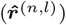. *Medium-sized* axon radii remained unchanged. *Small* and *large* axon radii were corrected (green) from annotated data. To generate the erroneous axon radii ensembles (b-e), either corrections were not applied (b,c), axon radii were manipulated (red; d) or a combination of both was applied (e). Note, that multiple (*M* = 500) axon radii ensembles were generated to assess the error introduced in the *medium-sized* and entire axon radii range, of which 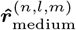 and 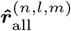 denote the *m*-th (*m ∈* [1, *M*]) axon radii ensemble, respectively.

*Generation of* 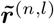. 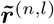 was generated by correcting the predicted axon radii ensemble 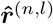 obtained from 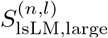 (see Table 2) in the range of *small* and *large* axon radii, whereas *medium-sized* axon radii remained unchanged (see Fig. 4a).

*Small* axon radii of 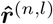 were corrected by substitution with the EM-based axon radii ensemble 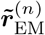 obtained from the *matching*, annotated subsection 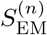 (see Table 2). As 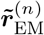 was smaller than 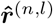, direct substitution was inappropriate. Instead, we discarded *small* axons of 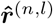 and randomly sampled *small* axons from 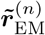 with replacement until the ratio of *small* and remaining axons equaled the ratio observed in 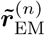.

*Large* axon radii of 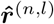 were corrected by substitution with the axon radii ensemble 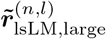 obtained from annotations on 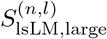 (see Table 2). To replace under- or oversegmented axons in 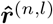 and to complement 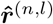 by falsely missed axons, we added axon radii of 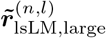. If existent, we replaced corresponding axon radii of 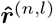 with a counterpart from 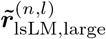. To determine a corresponding axon in 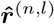 for each axon in 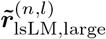, we computed the pairwise dice similarity as explained in Fig. 3. Remaining, *large* axon radii of 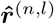 were considered falsely detected and were discarded for 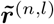.

*Generation of* 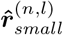. 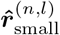 was generated analogously to 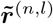 except that *small* axons were not corrected (see Fig. 4b). To estimate an upper bound of the error introduced by missing *small* axons, we repeated the experiment after discarding all *small* axons of 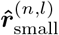.

*Generation of* 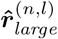. 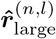 was generated analogously to 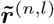 except that *large* axons were not corrected (see Fig. 4c).

*Generation of* 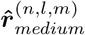. Due to the lack of annotated data for correction of *medium-sized* axons, we effectively used *medium-sized* axon radii of 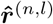 as a reference, i.e., 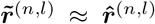 for *medium-sized* axons (see Fig. 4d). Erroneous, *medium-sized* axon radii were generated by manipulating 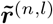 according to an error model in a Monte Carlo simulation (see Fig. 5). As a consequence of employing a Monte Carlo simulation approach, we did not generate one, but *M* = 500 erroneous axon radii ensembles. The parameters of the error model used in the Monte Carlo simulation were obtained from corresponding predictions and references (here: annotations) for the set of *N · L* = 30 subsections 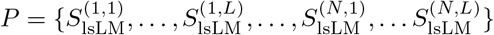 (see Table 2).

**Figure 5:**
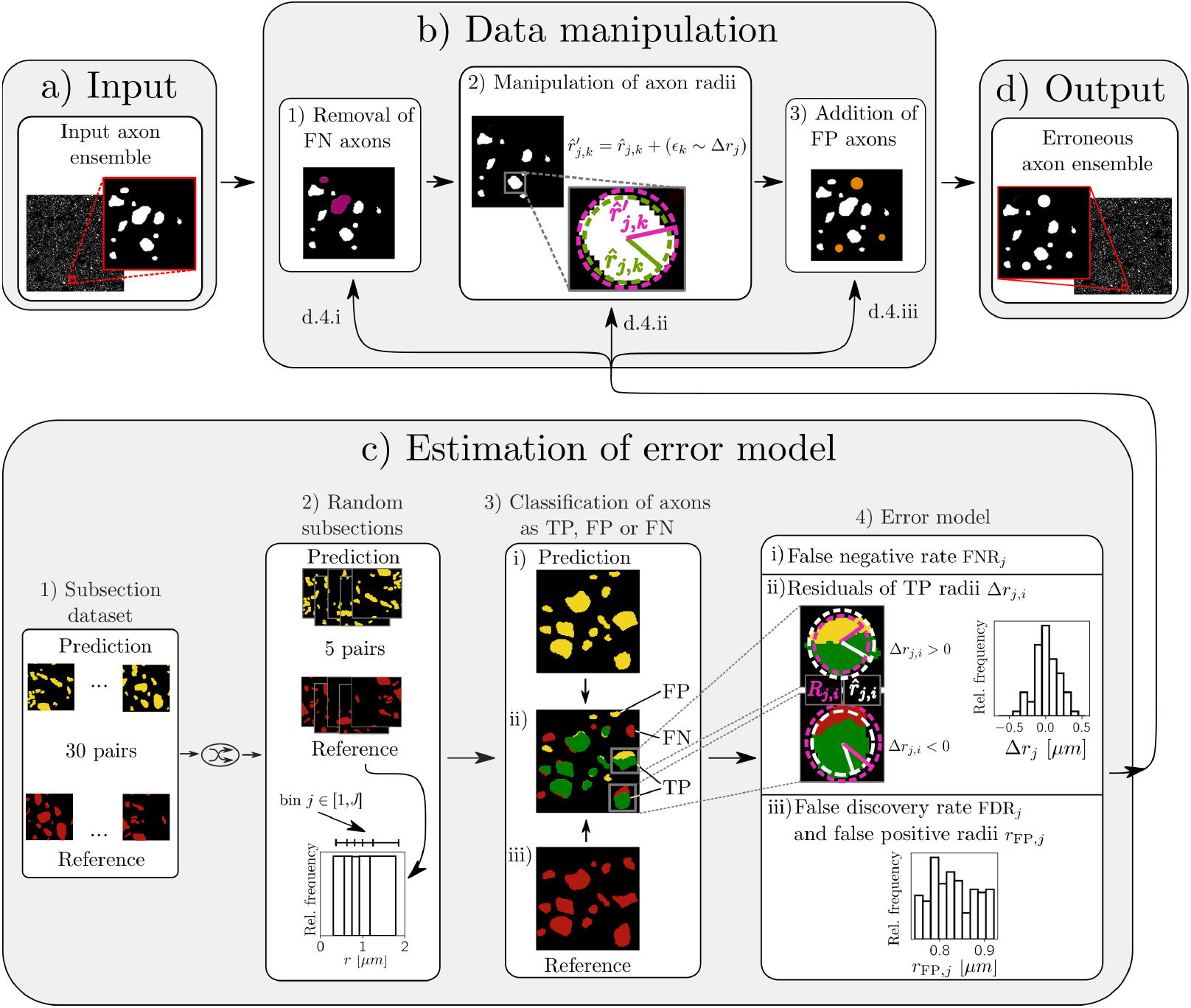
Schematic of the generation of axon ensembles with erroneous, *medium-sized* axon radii. The input axon ensemble (a) was manipulated (b) based on an error model (c) to generate axon radii ensembles with erroneous medium-sized axon radii (d). To model the error as a function of the axon radius, distinct parameters of the error model were determined per axon radii bin *j ∈* [1, *J* = 5] (see details on the binning below). To obtain an erroneous axon radii ensemble (d), we employed the error model (c.4) for each bin *j* as follows: First, randomly drawn, missed (false negative; FN) axon radii were removed (b.1, purple) according to the false negative rate (FNR_*j*_; c.4.i, see Eq. (6)). Then, axon radii were perturbed (b.2) according to the residuals of partially or fully detected (true positive; TP) axons (Δ*r*_*j*_; c.4.ii, see Eq. (7)). Finally, randomly drawn, falsely detected (false positive; FP) axon radii were added (b.3, orange) according to the false discovery rate (FDR_*j*_; c.4.iii, see Eq. (8)) and the distribution of FP axon radii (*r*_FP,*j*_; c.4.iii). To determine the parameters of the error model, we pooled over five pairs of corresponding predictions (c.2, yellow) and references (c.2, red) randomly drawn from 30 pairs (c.1). The *J* bins were chosen to contain the same number of reference axon radii per bin. For each bin *j*, axons of corresponding predictions (c.3.i) and references (c.3.iii) were compared (c.3.ii) to classify non-overlapping axons as FP (entirely yellow in c.3.ii) or FN (entirely red in c.3.ii) and partially or fully overlapping axons as TP (partially or fully green in c.3.ii and c.4.ii). Then (c.4), we determined the parameters of the error model.

To generate the *m*-th erroneous axon radii ensemble 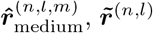 was manipulated as follows: first, randomly drawn, missed (false negative; FN) axons were removed; second, axon radii of correctly detected (true positive; TP) axons were perturbed; third, randomly drawn, falsely detected (false positive; FP) axons were added. The following paragraphs explain for the *m*-th iteration how model parameters were determined and how the so-obtained model was applied to generate 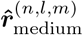. For brevity, we omit subsection, region and simulation iteration indices (*l, n* and *m*).

To determine the parameters of the error model, we pooled over five pairs of corresponding predictions and references for subsections randomly drawn from the aforementioned *P* (see Fig. 5c.1 and Fig. 5c.2). The error was modeled as a function of the axon radius. For this, we binned reference axons according to their radii into *J* = 5 bins, chosen to contain the same number of axons (see Fig. 5c.2, bottom). For each bin *j ∈* [1, *J*], corresponding predictions and references were compared to classify non-overlapping axons as FP or FN and fully or partially overlapping axons as TP (see Fig. 5c.3). Then, we counted the number of FN (*N*_FN,*j*_), FP (*N*_FP,*j*_) and TP (*N*_TP,*j*_) axons. Finally, we determined the false negative rate

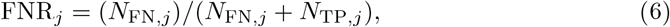

the residuals

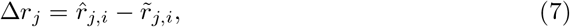

between predicted (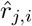 with *i* ∈ [1, *N*_TP,*j*_]) and reference (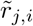 with *i* ∈ [1, *N*_TP,*j*_]) radii of TP axons, the false discovery rate

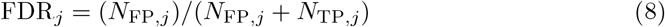

and the distribution of FP axon radii (*r*_FP,j_) (see Fig. 5c.4).

To apply the error model for bin *j*, the original axon radii ensemble with *N*_*j*_ axons in bin *j* was manipulated in three steps: First, 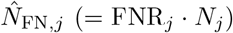 randomly drawn FN axons were removed (see Fig. 5b.1). Then, an error (*ϵ*_*k*_ with 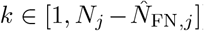) sampled from Δ*r*_*j*_ was added to each axon radius (see Fig. 5b.2). Finally, 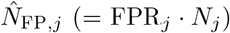 axons were added from the distribution of FP axon radii (*r*_FP,j_) to obtain the erroneous axon radii ensemble (see Fig. 5b.3 and see Fig. 5d).

*Generation of* 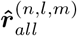. 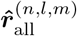 was generated analogously to 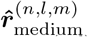 but with the following difference: instead of manipulating 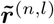, we manipulated 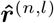 (see Fig. 4e).

From reference and erroneous axon radii ensembles, we computed effective axon radii 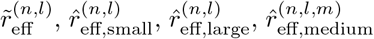 and 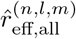 according to Eq. (3) for all *L*_large_ subsections of all *N* regions and all *M* simulation iterations. For 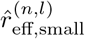 and 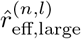, accuracy and bias with respect to 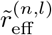 were assessed across all *N · L*_large_ subsections of all regions in terms of NRMSE and NMBE (defined analogously to Eqs. (4) and (5)). For 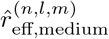 and 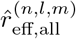, NRMSE_*m*_ and NMBE_*m*_ with respect to 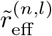 were computed per iteration *m* and summarized by their mean and standard deviation.

#### 2.6.4. Sensitivity of 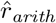 *and* 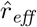 to variation of the image intensity

We assessed whether the influence of spatially varying intensity, e.g. introduced by staining heterogeneity, affected the capability of our method to map anatomy-related, spatial variation of 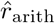 and 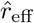 across whole lsLM sections. For qualitative analysis, we generated spatially smoothed maps of 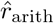 and 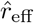 by computing the average of randomly positioned subsections (area: 0.12 µm^2^) and visually compared the patterns of the spatially smoothed maps to those of the corresponding lsLM images. For quantitative analysis, maps of 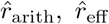 and the image intensity were generated similar to those above but sampled on an equally spaced grid (grid pixel area: 0.12 µm^2^). To obtain a scalar value for the image intensity, we applied gray scale conversion. Then, grid pixels of sections with similar axon radii distribution (G1, G2, M1, M2) were pooled and the correlation between image intensity and mapped radii was computed. As visual inspection suggested that *small* axons were particularly difficult to resolve in strongly stained areas, the above experiments were performed with and without considering *small* axons to test this hypothesis.

#### 2.6.5. Sensitivity of r_eff_ to outstandingly large axons

To evaluate how much *r*_eff_ is affected by outstandingly large axons, we investigated how *r*_eff_ changed as a function of a varying threshold *τ* when only axons with *r < τ* were considered for the computation of *r*_eff_. *τ* was chosen to cover the whole range of observed axon radii for a given ensemble. In particular, we assessed the worst case in which the largest individual axon was missed. To exclude estimation errors from this experiment, we considered only reference data, i.e., the reference axon ensembles generated in Section 2.6.3. Furthermore, to carry out this analysis at a scale as close as possible to the cross-sectional size of typical voxels of a human MRI system (1 mm^2^ or larger), we computed 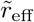 from combined axon radii ensembles for which we pooled over all *L*_large_ subsections per test region, yielding 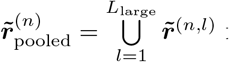 for the *n*-th test region. Thereby, we obtained 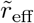 from the largest axon ensembles available for each test region based on combined areas of about 0.37 mm^2^.

## 3. Results

### 3.1. Performance of the semantic segmentation network

The segmentation metrics (balanced accuracy, dice, precision and recall, see Table 3) are given for all axons and only *large* axons. All metrics were higher when only *large* axons were evaluated than when all axons were evaluated.

**Table 3:** Segmentation metrics. Each *metric* was computed as the mean over all annotated subsections for *all axons* and only *large axons* (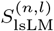 and 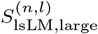; see Table 2).

| Metric | All axons | *Large* axons |
| --- | --- | --- |
| Balanced accuracy | 0.85 | 0.94 |
| Dice | 0.77 | 0.89 |
| Precision | 0.82 | 0.90 |
| Recall | 0.74 | 0.89 |

### 3.2. Error of 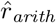

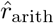 deviated from the line of unity, overestimating 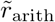 by 16 % (NMBE) and yielding an NRMSE of 21.5 % (see Fig. 6).

**Figure 6:**
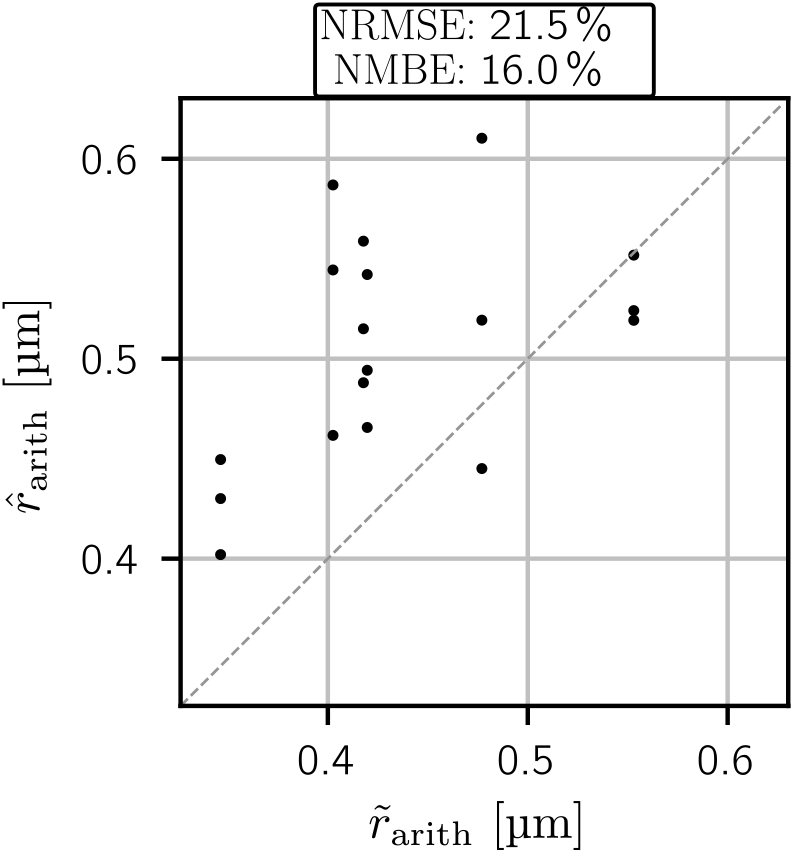
Error of 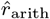. Each point compares an lsLM-based estimate 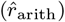 against its EM-based reference 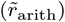. The dashed line represents the line of unity. NRMSE and NMBE over all subsections were 21.5 % and 16 %.

### 3.3. Error of 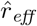

The overall accuracy and bias as assessed by NRMSE and NMBE were 7.2 % *±* 0.5 % and − 1.7 % *±* 0.4 % (see Fig. 7d). The agreement between 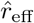 and 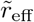 in the *small* (NRMSE: 0.6 % and NMBE: 0.4 %) and *medium-sized* (NRMSE: 1.9 % *±* 0.3 % and NMBE: 0.5 % *±* 0.4 %) axon radii range was higher than in the *large* (NRMSE: 7.4 % and NMBE: − 2.7 %) axon radii range (see Fig. 7a-c). The error in the *large* axon radii range translated into the range of all axons (see Fig. 7c-d). The error due to *small* axons increased when *small* axons were discarded for the estimation of 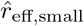 (NRMSE: 1.5 % and NMBE: 1.5 %) (see Fig. 7a).

**Figure 7:**
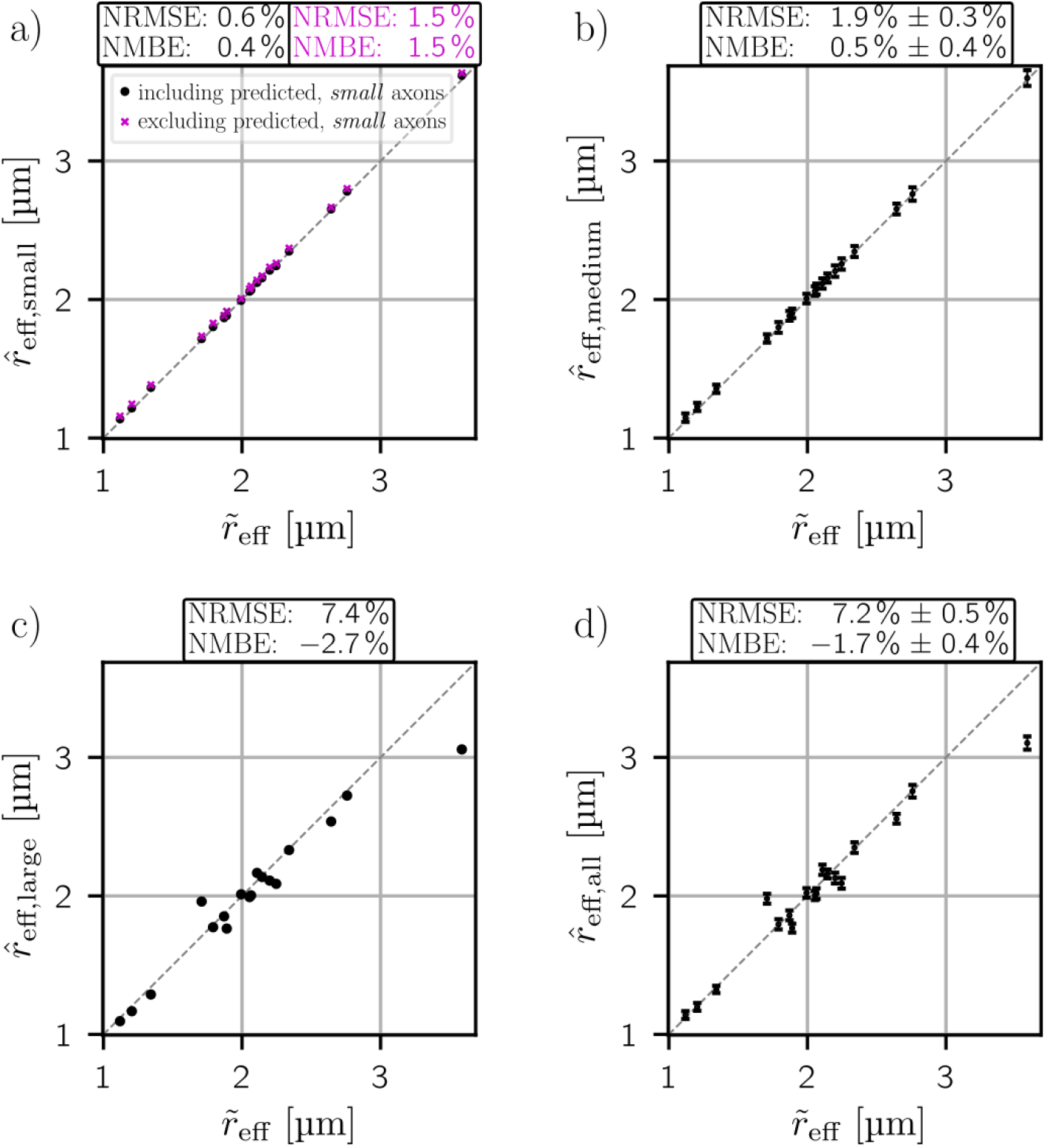
Error of 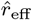. Depicted are comparisons of estimates 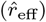 against a common reference 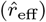, taking errors introduced in different axon radii ranges into account, i.e., *small* 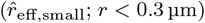 (a), *medium-sized* 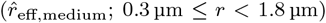 (b), *large* 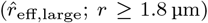 (c) and all 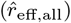 axons (d). Each point corresponds to a distinct lsLM subsection. The dashed line represents the line of unity. Two cases were considered for the error introduced in the *small* axon range: estimation of 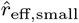 with (magenta) and without (black) including predicted, *small* axons. Error bars for results derived through Monte Carlo simulation (b, d) denote mean and standard deviation per subsection. NRMSE and NMBE were computed over all subsections according to Eqs. (4) and (5). Mean and standard deviation of NRMSE and NMBE across Monte Carlo simulation iterations are given, if applicable (b, d).

### 3.4. Sensitivity of 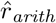 *and* 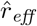 to variation of the image intensity

The spatial variation of *r*_arith_ resembled the image intensity distribution of the corresponding lsLM section (see Fig. 8a, top row and Fig. 8b). In contrast, maps of 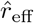 had a high local heterogeneity, which was not observed in the image intensity distribution of the corresponding lsLM section (see Fig. 8a, bottom row and Fig. 8b). These observations were supported by a strong correlation (*ρ* = 0.80, *p <* 10^*−*5^) between 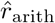 and the image intensity, which was reduced when *small* axons were discarded (*ρ* = 0.50, *p <* 10^*−*5^) (see Fig. 8c, top left and top right). In contrast, *r*_eff_ did not show a significant correlation with the image intensity (see Fig. 8c, bottom row).

**Figure 8:**
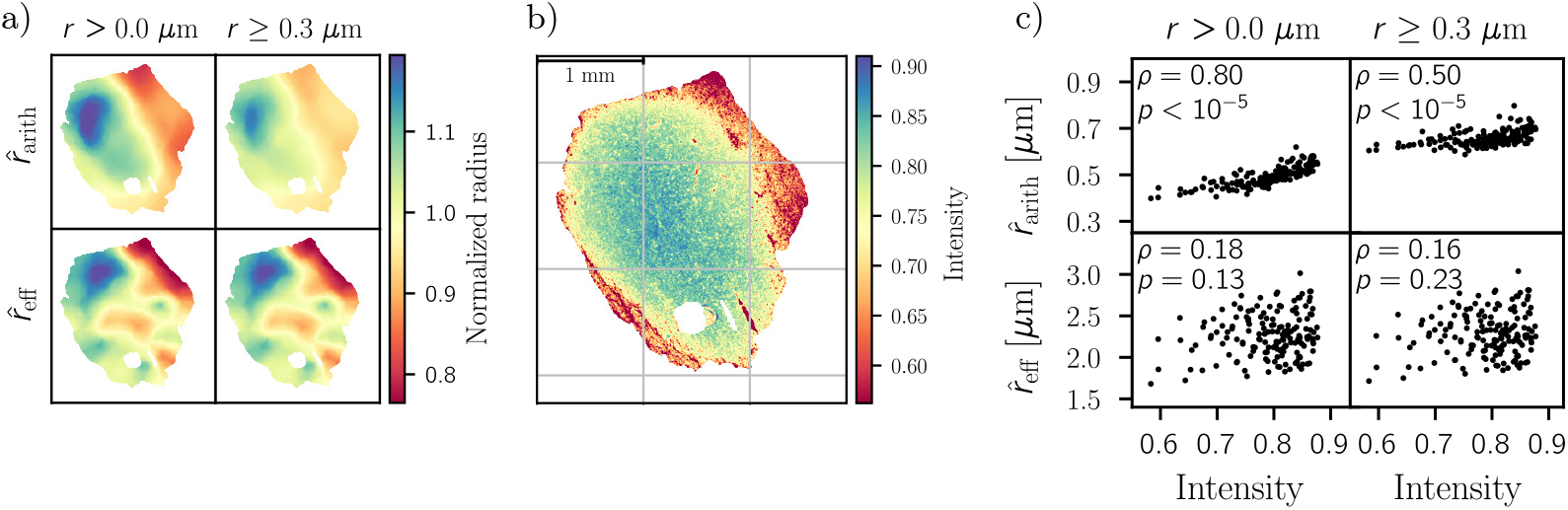
Sensitivity of 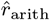 and 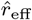 to variation of the image intensity. Depicted are: spatially smoothed maps of 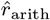 and 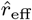 (a), the lsLM image of section M1 (b) adjusted to illustrate the correlation with maps of 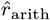 (a), and scatter plots between ensemble mean axon radii (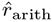 and 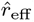) and lsLM image intensities (c). The correlation plots (c) pool across four sections (G1, G2, M1, M2). The *p*-values have been multiplied by the number of sections to correct for multiple comparisons (*ρ* is the correlation coefficient).

### 3.5. Sensitivity of r_eff_ to outstandingly large axons

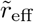 increased nonlinearly as a function of *τ* but with decreasing slope (see Fig. 9a). For large *τ*, the lines fragment due to the sparse occurrence of large axons. Compared to other regions, the influence of the largest axon on 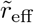 was particularly strong in region M2: 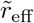 decreased by 14.5 % when the largest axon was discarded (see Fig. 9a). The largest axon was much larger other axons across all regions and its elongated shape suggested that this axon was oriented almost parallel to the cutting plane, i.e., its axon radius was strongly overestimated by the circular equivalent approximation (see Fig. 9b). The influence of the largest axon was smaller for the remaining regions: 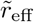 decreased by 0.9 % to 3 % when discarding the largest axon (see Fig. 9a).

**Figure 9:**
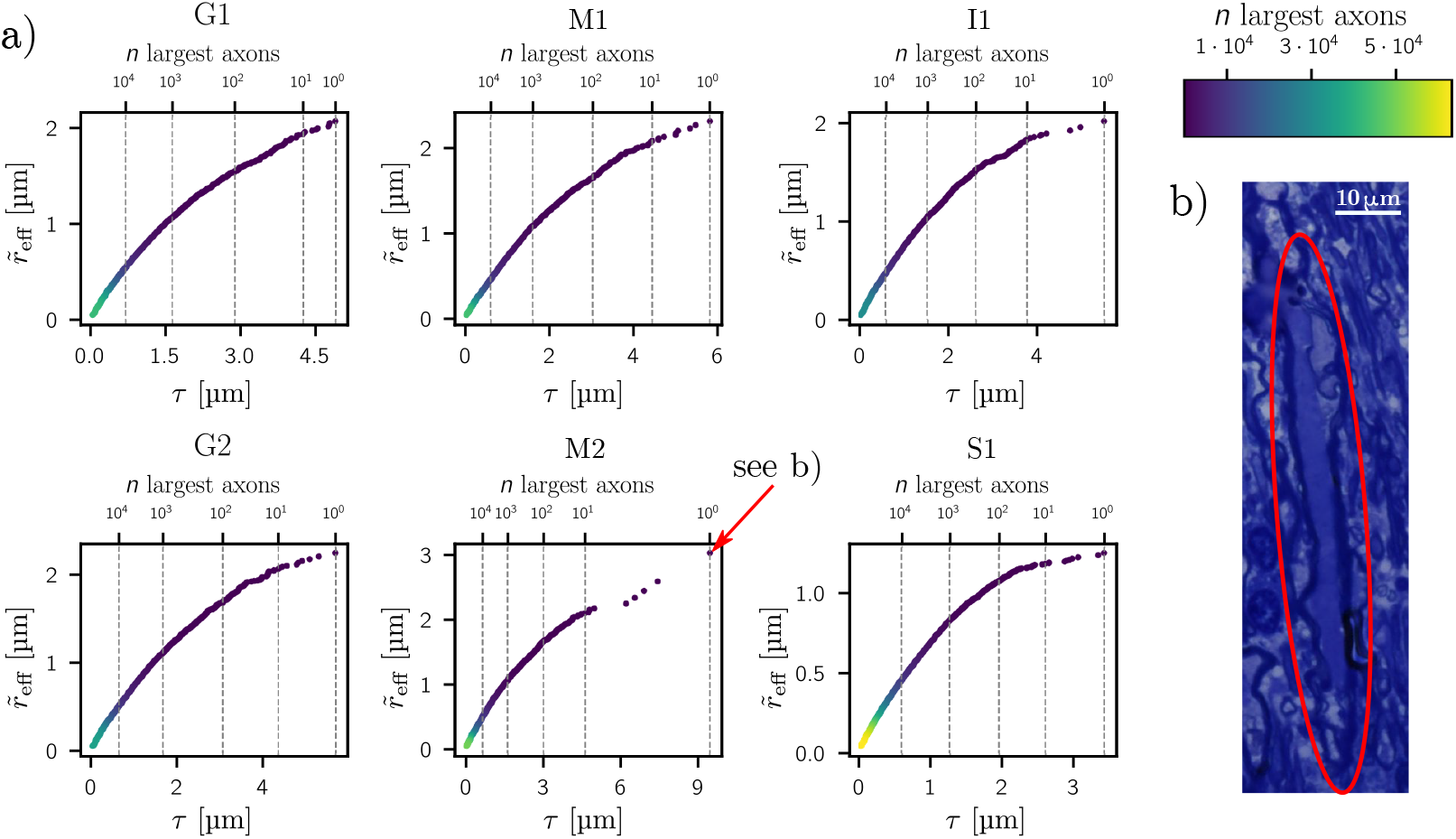
Sensitivity of *r*_eff_ to outstandingly large axons. Depicted are the values of 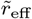 for the six test regions of the corpus callosum sample (a) when considering only axons with radius of *r < τ* for the computation of 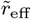. The marker colors indicate the order of axons sorted by their radius in descending order according to the colorbar in the top right. For orientation, powers of 10 are marked on top of the plots (a). The extracted lsLM subsection (b) shows the largest axon (*r* = 9.46 µm) observed across all regions, causing the jump of 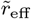 for region M2 between largest and second-largest axon. The elongated shape of this axon is likely due to the axon being oriented almost parallel to the cutting plane of the two-dimensional section. When discarding this axon, 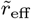 decreased from 3.02 µm to 2.58 µm, i.e., a decrease of 14.5 %. For the remaining regions, the decrease of 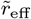 ranged from 0.9 % to 3 %. The total number of axons ranged from 3 *·* 10^4^ to 6.3 *·* 10^4^.

## Discussion

We investigated the potential of CNN-based segmentation on high-resolution, largescale light microscopy (lsLM) sections to narrow the scale gap between histological reference data and MRI voxels for the validation of MRI-based radius estimation in human brain tissue. The proposed pipeline accurately estimates the effective axon radius (*r*_eff_) in a human corpus callosum on sections spanning several cross-sections of typical voxels of human MRI systems (1 mm^2^ or larger) and is thus a promising candidate for the validation of MRI-based radius estimation in the human brain. However, the arithmetic mean radius (*r*_arith_), which is commonly reported in neuroanatomical studies, is less accurately estimated.

### Error of r_arith_ and r_eff_

To assess the estimation error of *r*_eff_ representatively for cross-sections of MRI voxels (1 mm^2^ or larger) of a human MRI system, sufficient sampling of the tail of the axon radii distribution is required. Therefore, we investigated large ensembles of axons including at least 10,000 axons per sample. To address the challenge of assessing the error of *r*_eff_ on large ensembles of axons, we divided the axon radii distribution into three ranges and employed distinct annotation and evaluation approaches. Thereby, we circumvented exhaustive annotation of smaller axons and enabled complete annotation of *large* (*r ≥*1.8 µm) axons, which we expected to be of greatest relevance for *r*_eff_.

Across the entire range of axon radii, we conclude higher suitability to estimate *r*_eff_ than *r*_arith_ due to higher accuracy (normalized-root-mean-square-error: 7.2 % vs. 21.5 %) and lower bias (normalized-mean-bias-error: −1.7 % vs. 16 %). Assessment of individual ranges revealed that erroneous, *large* axons predominantly determine the estimation accuracy of *r*_eff_. In contrast, erroneous, *small* axons (*r <* 0.3 µm) below the resolution limit of lsLM introduced only a minor overestimation, even when they were neglected altogether for estimating *r*_eff_. Thus, the potential of lsLM to sample the tail of the axon radii distribution in large field-of-views outweighs its limited capability to resolve *small* axons for mapping *r*_eff_.

While we assessed the presented pipeline with particular focus on the ensemble mean radii of segmented axons, i.e., *r*_arith_ and *r*_eff_, we employed pixel-wise optimization during training. The metrics commonly used to evaluate pixel-wise segmentation reflect the better suitability to estimate *r*_eff_: large axons, that determine *r*_eff_, were best segmented.

### Mapping anatomy-related, spatial variation across whole sections

Toluidine staining introduces low-frequency variation of the image intensity across whole lsLM sections. In a spatial correlation analysis, we identified this variation as a confounding factor for mapping *r*_arith_ but not for mapping *r*_eff_. In the light of moderate errors, spatial variation of *r*_eff_ seems anatomy-related. As *r*_arith_ was particularly confounded when *small* axons were taken into account, *small* axons seem particularly prone to staining effects. The inaccurate resolution of *small* axons may explain the observed overestimation of *r*_arith_. For *r*_eff_, the correlation with the image intensity was hardly affected by inclusion or rejection of *small* axons which underlines their minor contribution towards *r*_eff_.

### Sensitivity of r_eff_ to outstandingly large axons

Due to the tail-weighting of *r*_eff_, individual, outstandingly large axons may strongly contribute towards *r*_eff_ and thus strongly decrease estimation accuracy in case of erroneous segmentation. We assessed this potential source of error by discarding the largest axon for the computation of *r*_eff_ in axon ensembles of at least 30,000 axons.

The strongest contribution (14.5 % in region M2) of an individual axon was due to an outlier. Across the remaining regions, the contribution was smaller (0.9 % to 3 %), but still notable, considering that these axons represented only 0.002 % to 0.003 % of the axon ensembles. For the outlier-region M2, the largest axon (r = 9.46 µm) was oriented almost parallel to the cutting plane, resulting in an elongated shape. Thus, circular equivalent approximation may largely overestimate axon radii and bias the estimation of *r*_eff_. Instead, axon radii may be estimated based on the minor axes of ellipsoids fitted to the axon areas.

The investigated lsLM subsections (area: *∼*0.37 mm^2^) were smaller than the cross-section of a typical MRI voxel (1 mm^2^ or larger). In the latter, we expect reduced potential of individual axons to bias *r*_eff_ due to the larger axon ensemble size.

### Limitations and future directions

Although the method accurately estimated *r*_eff_ for different axon radii ensembles sampled across the corpus callosum, further investigation is required to assess how well the model generalizes and how well the overall method translates to other brain areas.

Recent, automated methods for large-scale axon segmentation use different acquisition techniques [24, 25] and segmentation algorithms [24, 25], which, however, were trained on perfusion-fixed brain tissue of mice or rats. Future studies can use our method as a benchmark to validate the aforementioned approaches for *r*_eff_ estimation in human brain tissue.

To assess *r*_arith_, we compared lsLM-based estimates against EM-based references from close-by cut sections. This choice of reference has two limitations: first, the EM-based axon ensembles were smaller, i.e., covering only 5 to 10 % of their lsLM-based counterparts; second, spatial misalignment arised from section-to-section distance and unknown in-section location. However, we assumed that representative estimation of the frequency-weighted *r*_arith_ is rather enabled by accurate resolution of frequently occurring axons than by large ensemble size or exact spatial alignment. Consequently, we regarded EM as a more suitable reference than lsLM because EM can resolve all frequently occurring axons, including *small* axons below the resolution limit of lsLM. Indeed, we found *small* axons to be particularly prone to variation of the image intensity in lsLM which in turn led to consistent overestimation of *r*_arith_. The overestimation of *r*_arith_ indicates that a bias of the method predominantly determined the inaccuracy of *r*_arith_ rather than inappropriate reference data.

The annotation of microscopy slides is prone to errors and inter-observer variability, in particular in the presence of staining and tissue degradation due to the immersion-fixation used in this study. Employing strategies that address noisy and uncertain annotations, e.g. by design of specific loss functions may improve axon segmentation accuracy [34] and thus radius estimation accuracy.

## Conclusion

The presented pipeline accurately maps the MRI-visible, effective radius (*r*_eff_) in a human corpus callosum sample by combining high-resolution, large-scale light microscopy (lsLM) with deep learning at the cross-sectional scale of typical MRI voxels (1 mm^3^ or larger). Since the pipeline is based on the fast, cheap and simple to perform lsLM measurement, it can easily be used beyond the realm of MRI-based radius models, e.g., to generate a representative, neuroanatomical atlas of the ensemble of large axons across the human corpus callosum. Generalization to other brain areas with different axon radii ensembles is yet to be demonstrated.

## List of Symbols and Acronyms

**Symbols**

**Axon radii ensemble**

r: individual axon radius
*r*: axon radii ensemble
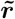: reference axon radii ensemble
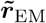: EM-based axon radii ensemble obtained through annotation of *S*_EM_
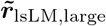: lsLM-based axon radii ensemble for *large* axons obtained through annotation of *S*_lsLM,large_
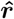: estimated axon radii ensemble
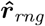: estimated axon radii ensemble with erroneous radii in axon radii range *rng*

**Axon radii ranges**

*small*: axons with *r <* 0.3 µm
*medium-sized*: axons with 0.3 µm *≤ r <* 1.8 µm
*large*: axons with *r ≥* 1.8 µm

**Arithmetic mean axon radius**

*r*_arith_: arithmetic mean radius
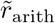: reference arithmetic mean radius
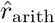: estimated arithmetic mean radius

**MRI-visible, effective axon radius**

*r*_eff_: effective radius
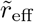: reference effective radius
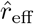: estimated effective radius
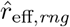: estimated effective radius based on an axon radii ensemble with erroneous radii in axon radii range *rng*

**Subsections**

*S*_EM_: EM subsection
*S*_lsLM_: small lsLM subsection
*S*_lsLM,large_: large lsLM subsection

**Acronyms**

CNN: convolutional neural network
EM: electron microscopy
lsLM: high-resoution, large-scale light microscopy
NMBE: normalized-mean-bias-error
NRMSE: normalized-root-mean-square-error

## Code and data availability

The source code and data used in this study will be made publicly available upon publication.

## Ethics

For samples used in this study, the entire procedure of case recruitment, acquisition of the patient’s personal data, the protocols and the informed consent forms, performing the autopsy and handling the autopsy material have been approved by the responsible authorities (Approval #205/17-ek).

## CRediT authorship contribution statement

**Laurin Mordhorst**: Conceptualization, Formal Analysis, Investigation, Methodology, Software, Visualization, Writing - Original Draft. **Maria Morozova**: Investigation, Writing - Review & Editing: **Sebastian Papazoglou**: Investigation, Supervision, Writing - Review & Editing. **Björn Fricke**: Investigation. **Jan Malte Oeschger**: Investigation, Writing - Review & Editing. **Thibault Tabarin**: Conceptualization, Investigation. **Henriette Rusch**: Investigation, Writing - Review & Editing. **Carsten Jäger**: Resources, Writing - Review & Editing. **Stefan Geyer**: Funding Acquisition. **Nikolaus Weiskopf**: Funding Acquisition, Writing - Review & Editing. **Markus Morawski**: Funding Acquisition, Resources, Writing - Review & Editing. **Siawoosh Mohammadi**: Conceptualization, Funding Acquisition, Methodology, Writing - Review & Editing, Supervision.

## Acknowledgements

The research leading to these results has received funding from the European Research Council under the European Union’s Seventh Framework Programme (FP7/2007-2013) / ERC grant agreement n° 616905.

This work was supported by the German Research Foundation (DFG Priority Program 2041 “Computational Connectomics”, [MO 2397/5-1; MO 2249/3–1; GE 2967/1-1], by the Emmy Noether Stipend: MO 2397/4-1) and by the BMBF (01EW1711A and B) in the framework of ERA-NET NEURON and the Forschungszentrums Medizintechnik Hamburg (fmthh; grant 01fmthh2017).

We are grateful to Dr. René Werner and Amra Hot for insightful discussions.

